# Gut-derived *Flavonifractor* species variants are differentially enriched during *in vitro* incubation with quercetin

**DOI:** 10.1101/2019.12.30.890848

**Authors:** Gina Paola Rodriguez-Castaño, Federico E. Rey, Alejandro Caro-Quintero, Alejandro Acosta-González

## Abstract

Flavonoids are a common component of the human diet with widely reported health-promoting properties. The gut microbiota transforms these compounds affecting the overall metabolic outcome of their consumption. Flavonoid-degrading bacteria are often studied in isolation under culture conditions that do not resemble the conditions in the colon and that eliminate the multiple interactions that take place in complex communities. In this study, a comparative metataxonomic analysis of fecal communities supplemented with the flavonoid quercetin led us to identify a potential competitive exclusion interaction between two sequence variants related to the flavonoid-degrading species, *Flavonifractor plautii*, that belong to the same genus but different species. During incubation of fecal slurries with quercetin, the relative abundance of these two variants was inversely correlated; one variant, ASV_65f4, increased in relative abundance in half of the libraries and the other variant, ASV_a45d, in the other half. This pattern was also observed with 6 additional fecal samples that were transplanted into germ-free mice fed two different diets. Mouse’s diet did not change the pattern of dominance of either variant, and initial relative abundances did not predict which one ended up dominating. Potential distinct metabolic capabilities of these two *Flavonifractor*-related species were evidenced, as only one variant, ASV_65f4, became consistently enriched in complex communities supplemented with acetate but no quercetin. Genomic comparison analysis of the close relatives of each variant revealed that ASV_65f4 may be an efficient ethanolamine-utilizing bacterium which may increase its fitness in media with no quercetin compared to ASV_a45d. Other discordant features between ASV_65f4- and ASV_a45d-related groups may be the presence of flagellar and galactose-utilization genes, respectively. Overall, we showed that the *Flavonifractor* genus harbors variants that present a pattern of negative co-occurrence and that may have different metabolic and structural traits, whether these differences affect the dynamic of quercetin degradation warrants further investigation.

## Introduction

Flavonoids are 3-ring phenolic compounds found in fruits and vegetables, their regular consumption is associated with health benefits (1,2). Among these, quercetin is one of the most abundant in the human diet. It exerts effects on the immune, digestive, endocrine, nervous, and cardiovascular systems (3–6). Some gut bacteria can cleave the central ring in the flavonoid skeleton by a process known as C-ring fission which generates smaller phenolic products. In the case of quercetin, phloroglucinol and 3,4-dihydroxyphenylacetic acid (DOPAC) are formed (7). It is not clear to what extent do the health effects of flavonoids depend on their transformation to biologically active compounds by the gut microbiota. Vissiennon and collaborators showed that the anxiolytic activity of quercetin is induced by DOPAC and not by the parent compound, evidencing a case in which the microbial metabolite exerts the beneficial effect (8). DOPAC has also antiproliferative activity in colon cancer cells (9) and anti-platelet aggregation activity (10). Additionally, the well-recognized quercetin-degraders, *F. plautii* (formerly *Clostridium orbiscindens*) and *E. ramulus*, are also butyrate-producers, a short-chain fatty acid that is the preferred source of energy of colonocytes and essential for colon health (11–14).

Most commonly, these flavonoid-degrading bacteria are studied in pure culture, however, they reside in the colon where more than 10^13^ bacterial cells inhabit (15). *In vitro* fecal incubation systems provide a scenario where complex microbial communities can be studied. *In vitro* fecal incubations with quercetin have revealed the metabolites formed during the transformation of this flavonoid; however, ecological interactions between quercetin-degraders and the rest of the community have been overlooked. Additionally, most studies have analyzed fecal samples from one donor (16,17) or pooled fecal samples from different donors (18–20), dismissing the importance of the gut microbiota at the individual level. To characterize key players in quercetin degradation that are common or unique among subjects, the evaluation of *in vitro* fecal incubation experiments with individual samples is needed.

In this study, microbial community dynamics from different subjects were analyzed individually by *in vitro* incubations of feces with quercetin. Through this approach, we identified variants related to flavonoid degraders that became enriched upon incubation with quercetin and that were consistently observed across 15 healthy subjects. Since it is known that differences in a small set of genes between bacterial strains can have a profound impact on the host’s physiology (21), we used a genomic comparison analysis of close relatives of these variants to infer genetic differences between them. Potential metabolic and structural differences were detected between these variants, thus, we propose that their role in the gut microbiome is differentially affected by carbon sources and interactions with other members of the community and may have distinct roles in the degradation of the flavonoid.

## Materials and Methods

### Sample collection and processing

Stool samples were collected from subjects participating in the Wisconsin Longitudinal Study (WLS) between November 2014 and February 2015 as previously described (22). Briefly, participants collected stool samples directly in sterile containers, then samples were kept at ∼4 °C until arrival to the processing laboratory within 48 hours of collection. Upon arrival, sterile straws were filled with the fecal material and stored at - 80° C as previously described (23). The use of WLS fecal microbiota was approved by the Institutional Review Board at the University of Wisconsin-Madison.

### *In vitro* incubations with human fecal samples and quercetin

Human fecal samples were used for *in vitro* incubations with quercetin. Samples from 9 subjects kept in frozen straws were aliquoted (∼50 mg) on dry ice, weighted, and resuspended at 0.5 mg ml^-1^ final concentration with heavy vortexing in anaerobic 7N minimal medium supplemented with 20 mM sodium acetate (filter-sterilized through a 0.22-m pore diameter) (24). Quercetin dihydrate, 97 % w/w (Alfa Aesar) concentration was 0.125 mg ml^-1^ (0.4 mM) in the corresponding medium. Controls consisted of the same medium plus fecal sample. Treatments with quercetin had three replicates and controls one replicate. Anaerobic bottles were kept statically at 37° C. When quercetin disappearance was visible, 10 % of the culture was transferred to another anaerobic bottle with the same medium. Sample processing was done in an anaerobic chamber under an atmosphere of nitrogen (75%), carbon dioxide (20%), and hydrogen (5%). Sampling was done at the end of the first and second incubation time once quercetin degradation was completed across all samples (72 h of incubation). In a second experiment, combinations of fecal matter from different subjects were tested. Two fecal samples used in the previous *in vitro* incubation experiment (from subjects #3 and #4) were selected based on their enrichment of one or the other *Flavonifractor*-related variant (ASV_65f4 or ASV_a45d). Three enrichments of these fecal samples were done as follows (#3/#4): 0/0.1, 0.1/0.1, 0.1/0 mg ml^-1^. Incubations were done again in anaerobic 7N minimal medium supplemented with 20 mM sodium acetate with and without quercetin using three replicates. Sampling was done at 0 and 72 h of microbial growth.

### *In vitro* incubations with fecal samples from human microbiota-associated mice (HMAM) under different diets

Experiments involving mice were performed using protocols approved by the University of Wisconsin-Madison Animal Care and Use Committee. Six female C57BL/6 (B6) germ-free mice were gavaged with ∼200 µl of fecal slurry which was prepared under anaerobic conditions in Hungate tubes using a 1 cm piece of straw containing the frozen fecal material and 5 ml mega media (25). Mice were maintained on a chow diet for 2 weeks after humanization, then, a diet high in fiber (Teklad 2018S) was administered for two weeks, and then switched to a low fiber diet for two weeks (Teklad TD.97184). Three fecal pellets were collected for each mouse after each experimental diet (high and low in fiber) and used separately as inoculum for *in vitro* incubations. Fecal pellets were weighted and then resuspended at 0.15 mg ml^-1^ final concentration with vigorous vortexing in anaerobic 7N minimal medium plus acetate and quercetin as described above. Sample processing and incubation were also done under anaerobic conditions at 37° C statically. Sampling was done at 0 and 7 days of incubation (only one incubation time was done).

### HPLC analyses of quercetin and metabolites

Samples were processed as previously described (24). Briefly, 200 μl samples were mixed with 1000 μl HPLC-grade methanol plus 20 μM genistein as internal standard, the suspension was bead beaten (2 min), heated (56 °C for 20 min) and spun (10 min at 18, 000 *g*). Then 1 ml of the supernatant was mixed with 200 μl of 10 mM ammonium formate/0.5 M EDTA buffer (pH 3.5). Separations were performed on a Kinetex 5 μm EVO C18, 100 Å, 250 × 4.6 mm column (Phenomenex, Torrance, CA, USA). Injection volumes were 10 µL. The flow rate was 1 ml min^-1^. Run time was 59 min run. The mobile phase was a binary gradient of (A) 10 mM ammonium formate and 0.3 mM ethylenediaminetetraacetic acid in water adjusted to pH 3.5 using concentrated HCl and (B) methanol. The gradient was: 5 % B for 5 min, increased to 30 % B over 30 min, increased to 95 % B over 10 min, remained constant at 95 % B for 5 min, decreased to 5 % B over 2 min, and then re-equilibrated at 5 % B for 7 min. Chromatograms at 280 nm absorbance were analyzed.

### DNA preparation

A 300 µL aliquot of each culture was mixed with a solution containing 500 µl of 200 mM Tris (pH 8.0), 200 mM NaCl, 20 mM EDTA, 200 µl of 20 % SDS, 500 µl of phenol:chloroform:isoamyl alcohol (25:24:1, pH 7.9) and 1.2 mg of 0.1-mm diameter zirconia/silica beads (BioSpecProducts). The suspension was bead beaten (3 min), spun at 8,000 rpm (5 min), and then top layer was transferred to a 15 ml tube for immediate column purification with 2.5 vol of NTl buffer, 3 washes with NT3 and final elution with 25 μl of elution buffer (Clontech, Machery-Nagel 740609.250).

### 16S rRNA gene V4 amplification and sequencing

PCR was performed using primers 515F and 806R for the variable 4 (V4) region of the bacterial 16S rRNA gene (26). PCR reactions contained 1 ng µl^-1^ DNA, 10 µM each primer, 12.5 µl 2X HotStart ReadyMix (KAPA Biosystems, Wilmington, MA, USA), and water to 25 μl. PCR program was 95 °C for 3 min, then 30 cycles of 95 °C for 30 s, 55 °C for 30 s, and 72 °C for 30 s, the final step was 72 °C for 5 min. PCR products were purified by gel extraction from a 1.5 % low-melt agarose gel using a Zymoclean Gel DNA Recovery Kit (Zymo Research, Irvine, CA). Samples were quantified using the Qubit dsDNA HS assay (Invitrogen, Carlsbad, CA, USA) and equimolar concentrations pooled. The pool was sequenced with the MiSeq 2×250 v2 kit (Illumina, San Diego, CA, USA). All DNA sequences generated in this study are deposited in the NCBI Short Read Archive under BioProject Accession Number: PRJNA595821 and PRJNA596889.

### 16S rRNA sequence analysis

Sequences were demultiplexed on the Illumina MiSeq, and sequence clean-up was completed in Qiime2 Core 2018.11 (https://qiime2.org). Quality control, including removal of chimeras was performed with the DADA2 pipeline. The first 10 nucleotides were trimmed, and reads were truncated to 220 bases. DADA2 generates high-resolution tables of amplicon sequence variants (ASVs) which represent biological sequences in the sample differing by as little as one nucleotide (27). Taxonomy was assigned to ASVs using the feature-classifier classify-sklearn. The abundance of the resulting taxonomy assignments of ASVs was analyzed using STAMP 2.1.3 (statistical analysis of taxonomic and functional profiles) (28), with statistical comparisons between groups (e.x. control *vs*. quercetin treatment) performed by two-sided Welch’s t-test within 95% confidence interval. A subset of 6 ASVs whose abundance increased in quercetin treatments was further analyzed using SILVA ACT (Alignment, Classification and Tree Service) (29), and 10 closest neighbors were downloaded from this analysis. ASVs that lack neighbors with a defined taxonomy at the genus level in SILVA ACT were subjected to BLASTn (https://blast.ncbi.nlm.nih.gov/) using the Whole-genome shotgun contigs (WGS) database for Clostridia (taxid:186801), 16s rRNA partial sequences from the most similar genomes with a defined taxonomy at the genus level were used for the phylogenetic analysis. This group of sequences was aligned and analyzed in MEGA 6.06 (30), the alignment file was used to construct a phylogenetic tree using the UPGMA method and a Distance Matrix for estimating evolutionary divergence between sequences (30–32). Complete ids for ASV enriched in quercetin treatments and accession numbers for reference sequences are listed in Fig 1 (only the first 4 letters of each ASV are going to be mentioned throughout the text). Correlations (Spearman’s rs and Bonferroni correction) and Principal component analysis (PCA) were done using PAST 3.23 (PAleontological STatistics) (33).

**Fig 1.**
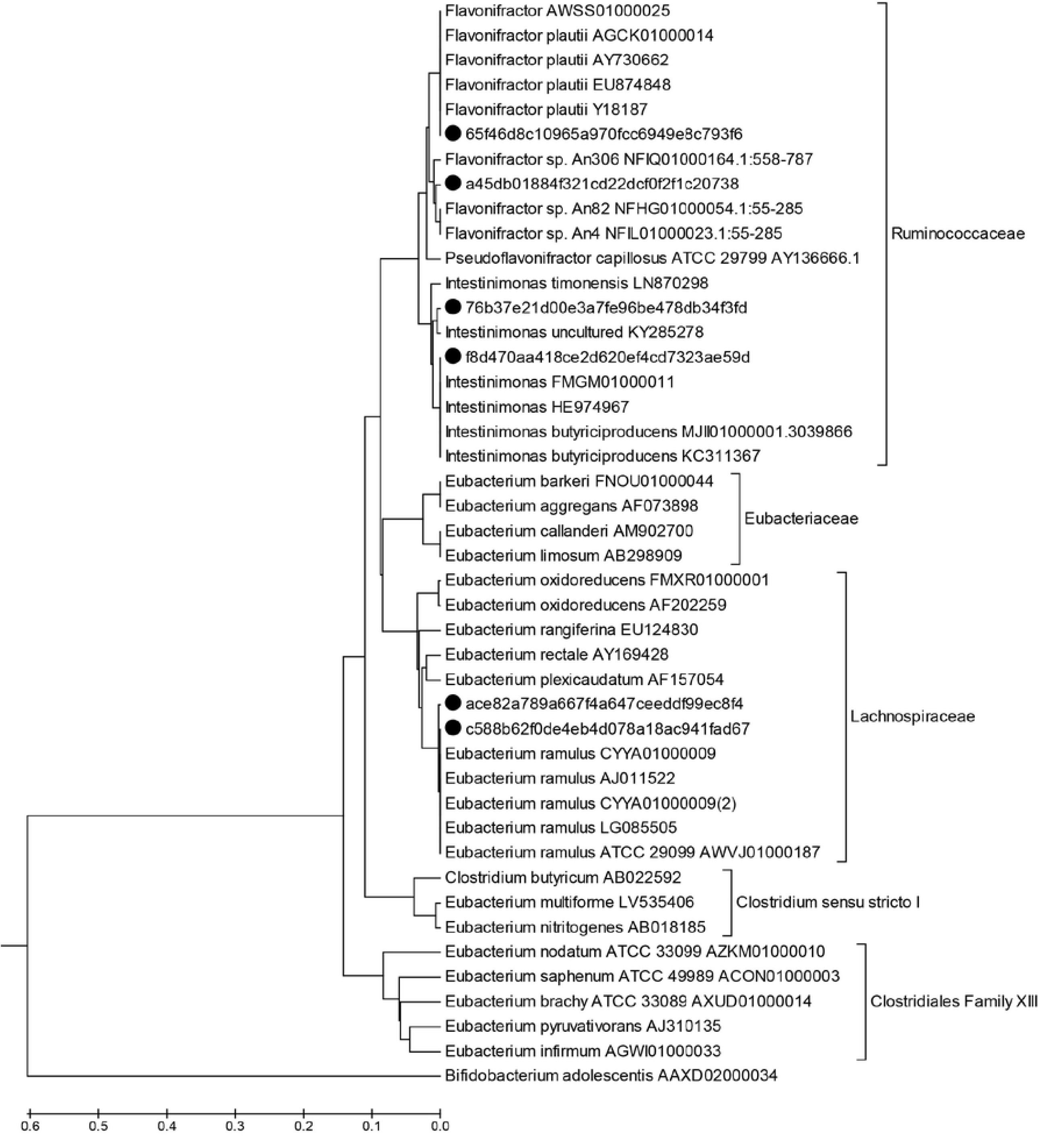
Phylogenetic analysis of taxa enriched in the presence of quercetin. The phylogenetic tree shows six ASVs (black dots) whose abundance increased in the presence of quercetin; distances in the tree were inferred using the UPGMA method (32). The optimal tree with the sum of branch length = 2.18101115 is shown. The tree is drawn to scale, with branch lengths in the same units as those of the evolutionary distances used to infer the phylogenetic tree. The evolutionary distances were computed using the Maximum Composite Likelihood method (31) and are in the units of the number of base substitutions per site. The analysis involved 44 nucleotide sequences. All positions containing gaps and missing data were eliminated. There were a total of 225 positions in the final dataset. Evolutionary analyses were conducted in MEGA6 (30).

### Selection of *Flavonifractor* variants-related genomes used in this study

Twenty genome assemblies for the genus *Flavonifractor* were downloaded from the NCBI genomes database in October 2019 (https://www.ncbi.nlm.nih.gov). Accession numbers and information about completeness is presented in S1 Table. To determine the species relationship of the genomes, the Average Nucleotide Identity (ANI) was calculated using the online tool JSpeciesWS (http://jspecies.ribohost.com/jspeciesws/) (34) which performs pairwise comparisons between two genomes calculating and indicating if a pair of genomes belong to the same species and/or genus based on their percentage of identity. After this analysis (S2 Table), we selected 8 genomes: 4 that represent the general features of the species *F. plautii* which is the closest relative of ASV_65f4, and 4 genomes which represent the general features of the genus *Flavonifractor*, including the ones most closely related to ASV_a45d (*Flavonifractor* sp. strains An4 and An82) (S1 and S2 Figs show the phylogenetic relatedness of *F. plautii* and *Flavonifractor* sp. strains, respectively).

### Genome comparative analysis

Annotation of functions was done using GhostKOALA (KEGG Orthology And Links Annotation, https://www.kegg.jp/ghostkoala/), an automatic annotation and mapping service using the database ‘genus_prokaryotes’ (35). Then prediction of orthologous gene clusters was done using OrthoVenn2 (https://orthovenn2.bioinfotoolkits.net/home) (36). We applied OrthoVenn2 clustering to identify gene clusters enriched in the groups most related to variant ASV_65f4 or ASV_a45d. Completeness of pathways was screened using the KEGG Mapper Reconstruction tool (https://www.genome.jp/kegg/tool/map_pathway.html). Proteins similar to a flavonoid-degrading protein was searched using the Blast tool in OrthoVenn2, Phloretin hydrolase (OXE48401.1) was taken as a reference protein (37).

## Results

### Taxa enriched in *in vitro* fecal incubations with quercetin belong to the Ruminococcaceae and Lachnospiraceae families

Microbial communities were monitored in two successive *in vitro* incubations of human fecal samples with and without quercetin. In all quercetin treatments, the main metabolite produced was DOPAC (S3 Table). Using STAMP statistical analysis, it was revealed that two groups of bacteria were significantly enriched in quercetin treatments; these were unidentified members of the Ruminococcaceae and Lachnospiraceae families (S3 Fig). We then determined significant differences in abundance profiles of individual Amplicon Sequence Variant (ASVs). Six ASVs were identified as being enriched in one or more libraries when comparing quercetin treatments *vs* controls (S4 Table), four belonged to the Ruminococcaceae family and two to the Lachnospiraceae. This group of sequences was subjected to a more detailed phylogenetic analysis which revealed that the closest relatives were members of the genera *Eubacterium* (Lachnospiraceae), *Flavonifractor* (Ruminococcaceae) and *Intestinimonas* (Ruminococcaceae) (Fig 1). As a result of this phylogenetic analysis, we identified ASVs that were 100 % identical to *F. plautii* (ASV_65f4) and *E. ramulus* (ASV_c588) (S5 Table). These ASVs were the ones that increased in abundance the most when quercetin was present compared to the controls, together with another one related to the *Flavonifractor* genus (ASV_a45d) (Fig 2). Although ASVs related to *Eubacterium* and *Intestinimonas* genera were enriched significantly in 1 or 2 libraries, *Flavonifractor*-related ASVs were found to be more ubiquitous after quercetin incubation, with at least one variant significantly enriched in every library (S4 Table). It should be noted that one of these *Flavonifractor*-related variants, ASV_65f4, showed also a significant increase when no quercetin was present in the medium (S4 A Fig), indicating that this ASV is favored by the culture conditions used. Nevertheless, its relative abundance increased significantly more when quercetin was present (S4 B Fig). This behavior did not change when higher concentrations of fecal matter were tested (1 and 10 mg/ml). The other *Flavonifractor* variant, ASV_a45d, showed no enrichment in media with no quercetin.

**Fig 2.**
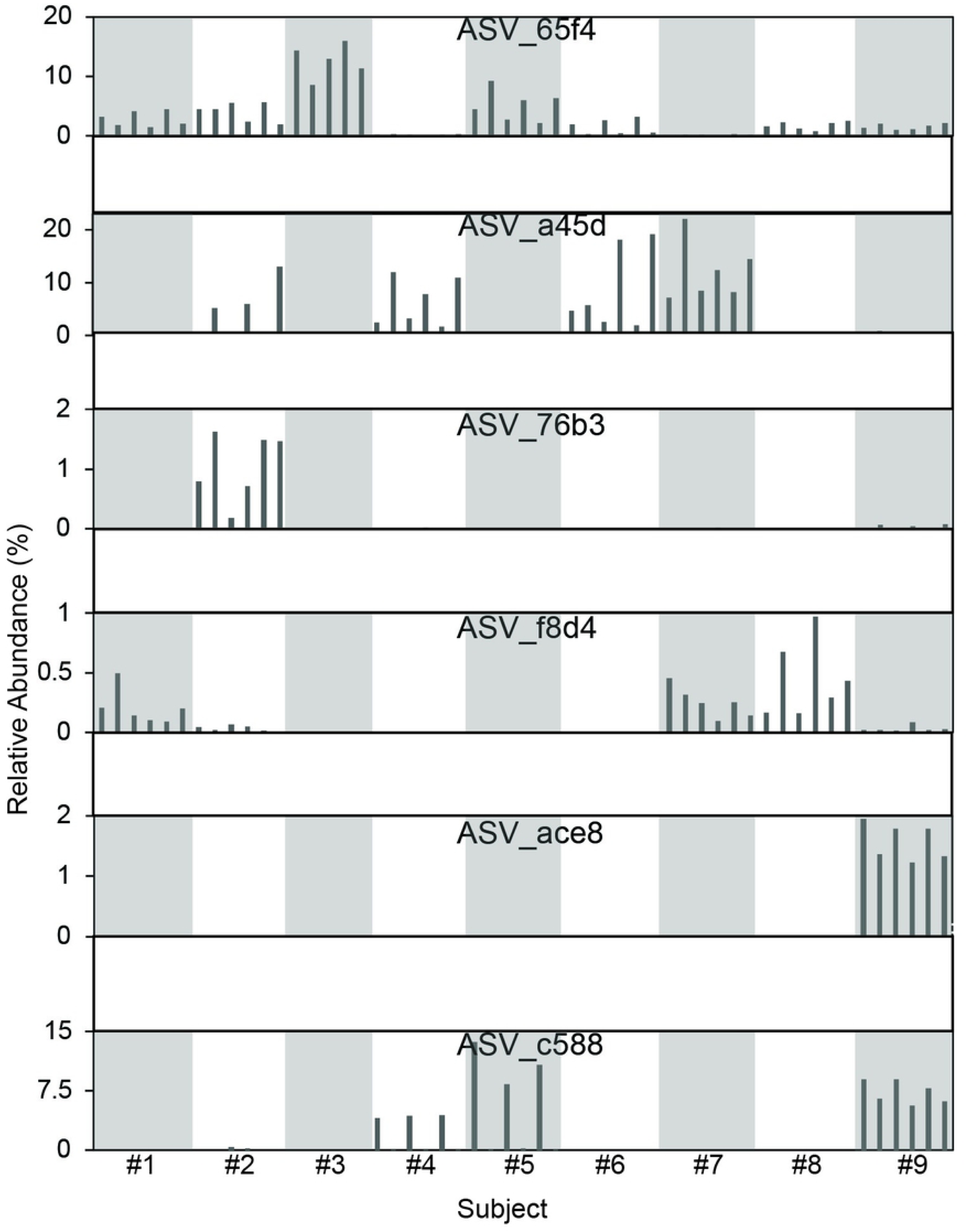
Bar plots for the relative abundances of the 6 Amplicon Sequence Variants (ASV) enriched in quercetin. Human subjects are labeled as Subject #1, #2, #3, #4, #5, #6, #7, #8, #9 in bar plots, for better visualization intercalated libraries are highlighted in gray. Each library has 6 bars, 3 corresponds to replicates from first incubation and 3 from second incubation (Subject #3 had only 2 replicates for second incubation).

### Amplicon Sequence Variants (ASV) related to *Flavonifractor* were negatively correlated in *in vitro* incubations with fecal samples

A correlation analysis across the 9 libraries of the abundances of the ASVs enriched in quercetin treatments showed a strong negative correlation between ASV_65f4 and ASV_a45d (Figs 3 A and S5 Fig), both of these ASVs were related to the genus *Flavonifractor* (Fig 1); this negative correlation was not present in incubations without quercetin. This pattern was investigated in a second experiment using human microbiota-associated mice (HMAM). Germ-free mice were inoculated with fecal samples from six human subjects different from the previous ones and after a period of acclimatization to the diet, fecal pellets were retrieved and used for *in vitro* incubations with quercetin. Both ASVs, ASV_65f4 and ASV_a45d, were present in all 6 subjects, and after quercetin treatment, a negative correlation between these two ASVs was again evident (Fig 3 B). Which ASV dominated during incubation with quercetin could not be explained by their initial abundances (S6 Fig). Even though ASV_a45d was lower in abundance than ASV_65f4 (undetectable in most cases), it dominated in half of the libraries after quercetin treatment. Additionally, mice were fed two different diets (high and low in fiber) which affected the composition of the microbial community as shown in a principal component analysis (PCA) (Fig 4 A). Nevertheless, this disturbance did not affect the pattern of dominance of ASV_65f4 over ASV_a45d or vice versa, previously observed (Fig 3 B). PCA analysis showed that component 2 was explained by diet at 0 days of incubation (Fig 4 A) and by the enrichment of the *Flavonifractor*-related variants (ASV_65F4 or ASV_a45d) at day 7 (Fig 4 B). *E. ramulus*-related ASVs (ASV_c588 and ASV_ace8) were not present in these libraries, and only one *Intestinomonas*-related ASV was present (ASV_f8d4), which increased in relative abundance during incubation with quercetin.

**Fig 3.**
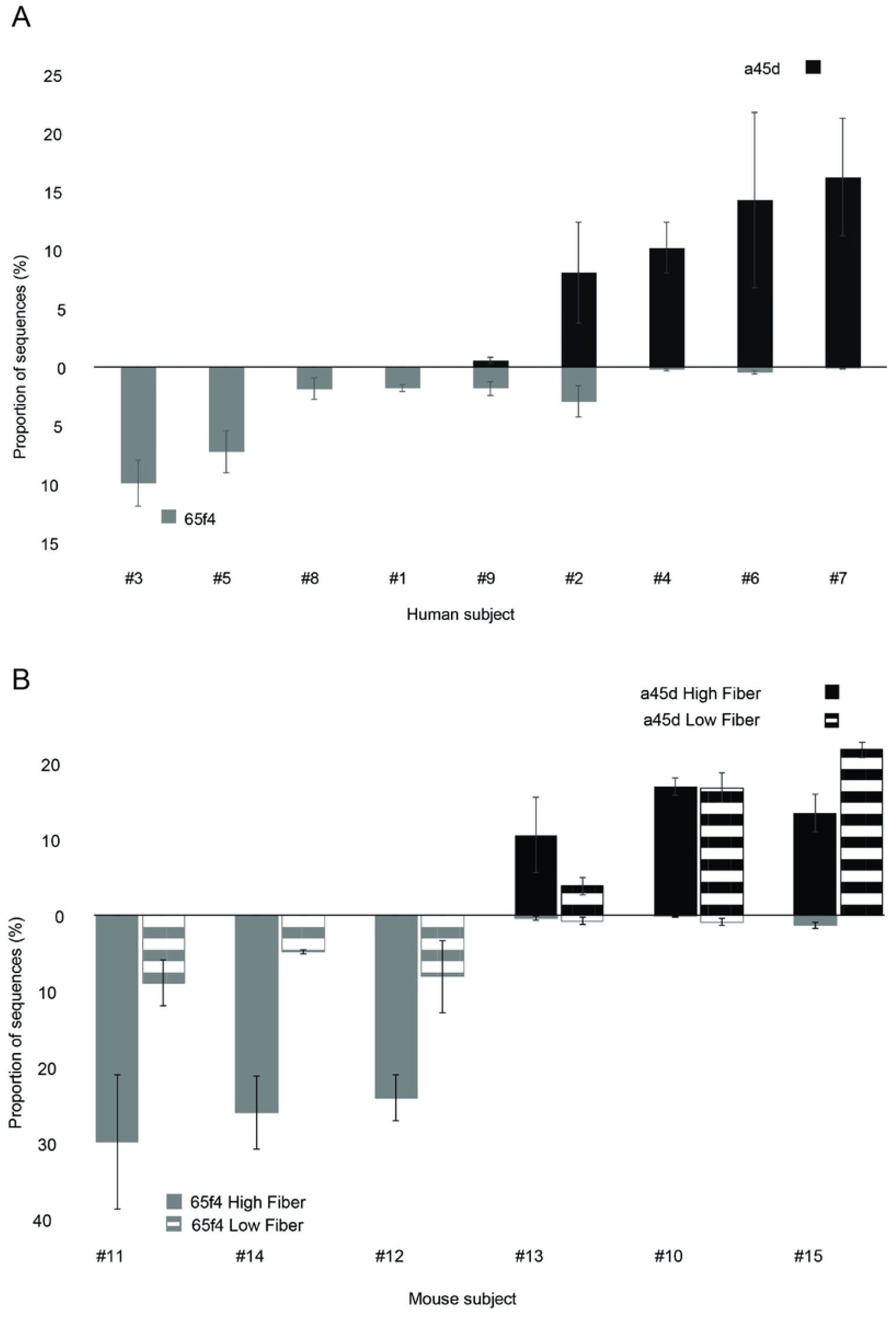
Relative abundances of ASV_65f4 and ASV_a45d are negatively correlated. Relative abundance of ASV_65f4 and ASV_a45d in human (A) and human microbiota-associated mice fecal samples (B). ASV_65f4 is represented in gray and ASV_a45d in black. In the bottom panel, libraries from HMAM mice fed a diet high in fiber are shown in solid color and mice fed a diet low in fiber are shown with a line pattern. Error bars correspond to 3 incubations done with fecal matter from the same donor individually sampled. For human samples (n=9), relative abundances obtained for the second incubation are shown and for HMAM mice (n=6), relative abundances obtained after 7 days of incubation with quercetin are shown.

**Fig 4.**
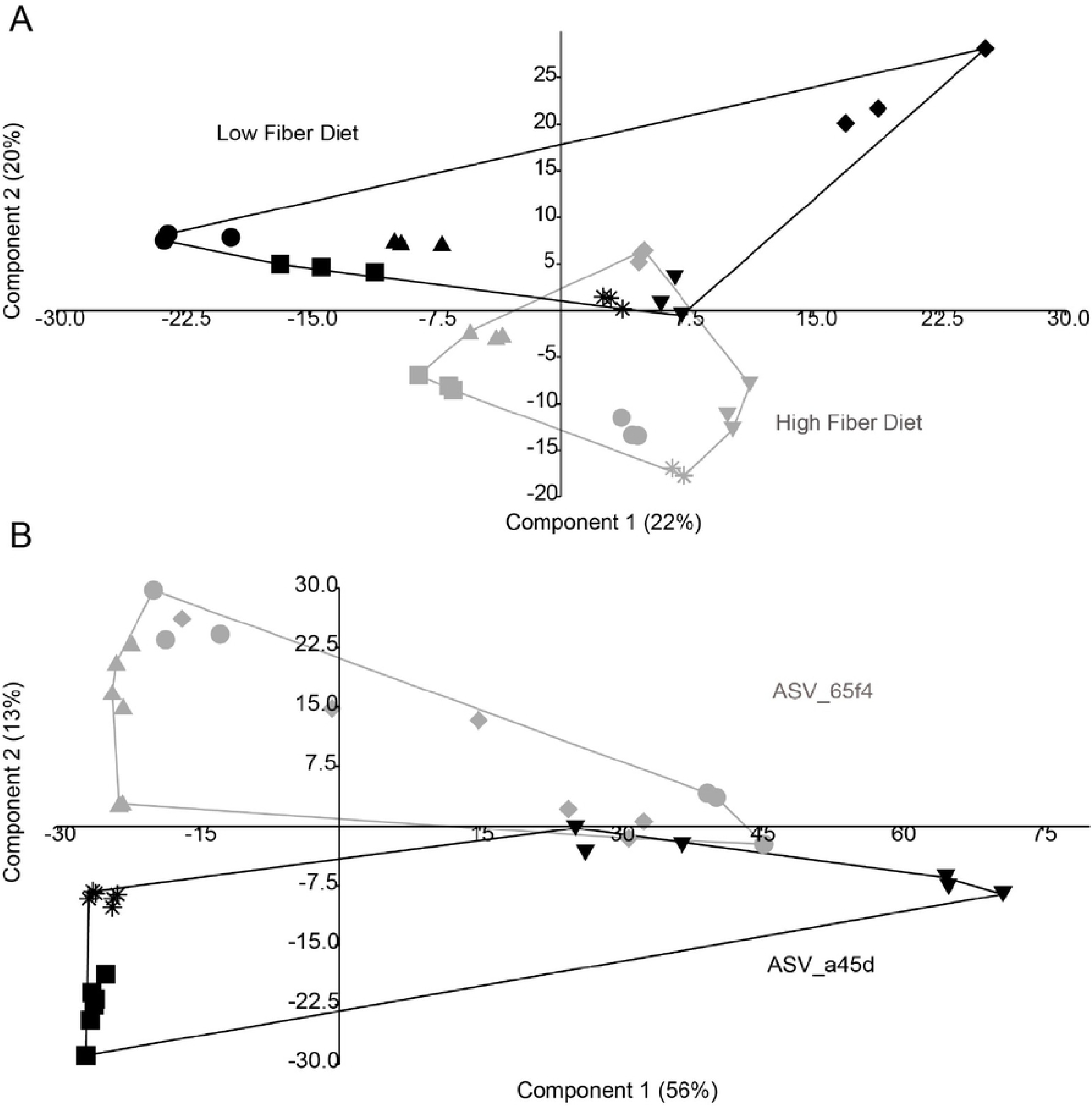
Principal component analysis (PCA) plot of the libraries from *in vitro* incubations with fecal samples from human microbiota-associated mice (HMAM). (A) PCA plot of the HMAM samples at 0 days of incubation. Libraries from HMAM mice fed a high fiber diet are shown in gray and libraries from HMAM mice fed a low fiber diet in black. (B) PCA plot of the HMAM samples at 7 days of incubation. Libraries that were enriched in ASV_65f4 are shown in gray and libraries enriched in ASV_a45d are shown in black. Each symbol represents a library (n=6, 3 replicates), different shapes represent libraries from a different subject: #10, star (*); #11, diamond (♦); #12, dot (●); #13, inv. triangle (▼); #14, triangle (▲); and #15, square (■).

### Fecal sample combinations showed the dominance of ASV_65f4 over ASV_a45d

After replicating the biological phenomenon between ASV_65f4 and ASV_a45d using fecal samples from different subjects, we aimed to study this pattern in cocultures. Unfortunately, only ASV_65f4 was isolated in pure culture (data not shown) while ASV_a45d could not be isolated. Thus, an experiment combining fecal samples that were previously enriched in ASV_65f4 or ASV_a45d was carried out (combination of fecal samples from subjects #3 and #4 previously enriched in ASV_65f4 and ASV_a45d, respectively, Fig 3 A). In this experiment, we expected that if the combination of fecal matters did not affect the *Flavonifractor*-related variants, we should observe a reduction in their relative abundance corresponding only to the dilution factor. This means that when samples #3 and #4 were combined, the relative abundances of ASV_65f4 and ASV_a45d should reach 50 % of the one reached when fecal samples are not combined. However, it was observed that when samples #3 and 4# were combined (50:50), ASV_65f4 dominated reaching a relative abundance compared to the one reached when fecal samples #3 and #4 were not mixed. Meanwhile, the relative abundance of ASV_a45d was severely affected by the combination of fecal samples reaching a relative abundance below 1 % (Fig 5).

**Fig 5.**
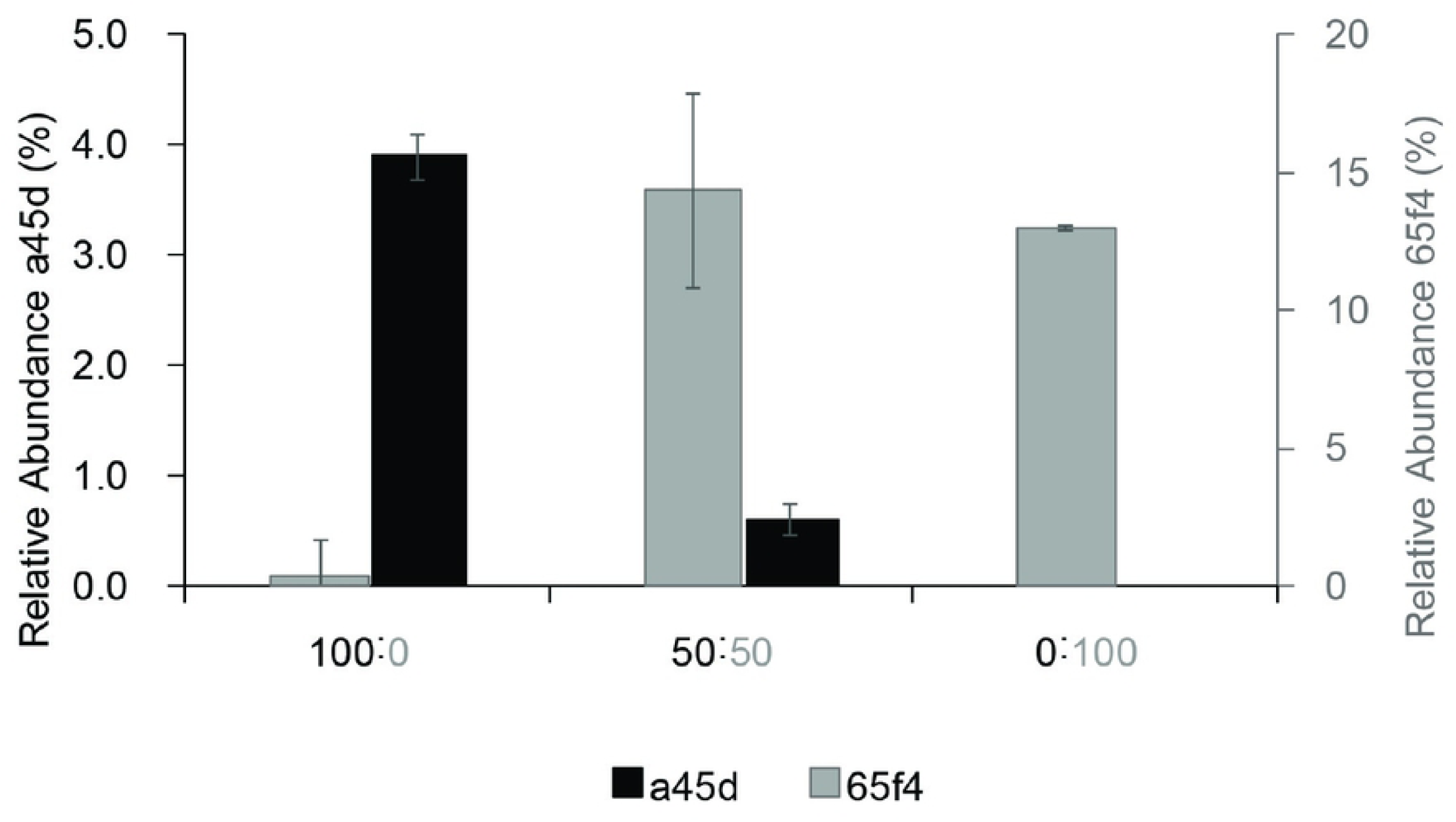
Relative abundance of ASV_65f4 and ASV_a45d in *in vitro* incubations with fecal sample combinations. Fecal samples from subjects #3 and #4 were combined (50:50) or not (100:0 and 0:100). *In vitro* incubations with the fecal samples from subject #3 (enriched in ASV_65f4) are shown in gray (right) and from subject #4 (enriched in ASV_a45d) is shown in black (left).

### Dominance of ASV_a45d or ASV_65f4 was associated with genera *Desulfovibrio* and *Phascolarctobacterium*

The microbial community profiles were analyzed in search of organisms whose relative abundance was lower or higher when ASV_a45d dominated over ASV_65f4 or vice-versa across all experiments. Besides, we only considered those species whose relative abundance increased during incubation because these species may have a higher probability of being actively interacting with other members of the community. It was observed that when ASV_a45d dominated after incubation with quercetin, the relative abundance of *Desulfovibrio* was significantly higher (p<0.05 in all 3 experiments) (Fig 6 A). When present, *Desulfovibrio* sp. increased in relative abundance in the medium supplemented with acetate. Meanwhile, when ASV_65f4 was dominant, the relative abundance of the genus *Phascolarctobacterium* was significantly higher (p<0.05 in all 3 experiments) (Fig 6 B). This genus increased in relative abundance during incubation as well.

**Fig 6.**
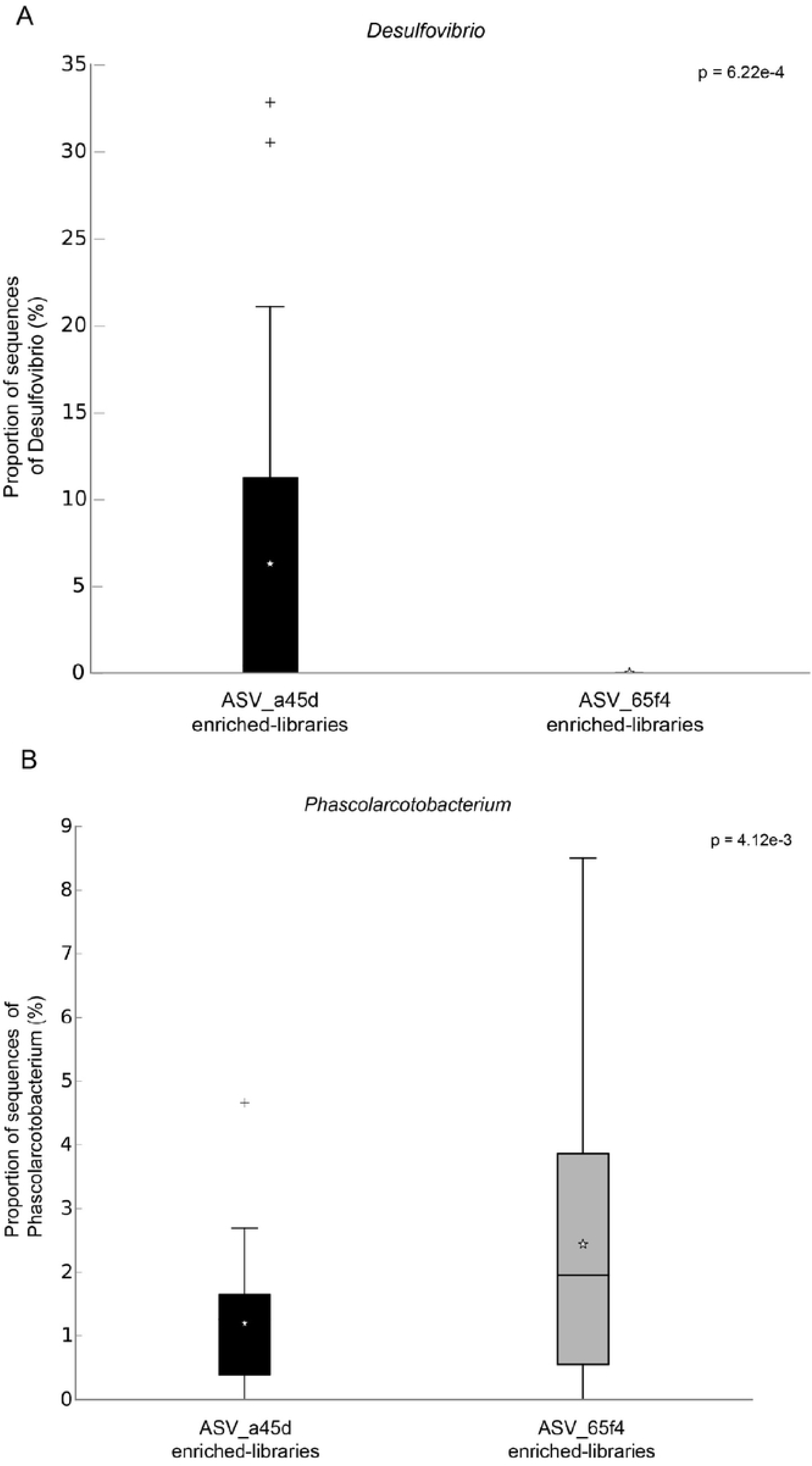
Box plots for the relative abundances of the genera *Desulfovibrio* and *Phascolarctobacterium.* (A) Box plots for the relative abundances of the genera *Desulfovibrio* in *in vitro* incubations with quercetin in libraries enriched in ASV_65f4 *vs* libraries enriched in ASV_a45d. (B) Box plots for the relative abundances of the genera *Phascolarctobacterium* in *in vitro* incubations with quercetin in libraries enriched in ASV_65f4 *vs* libraries enriched in ASV_a45d. Analysis for libraries from human subjects #1-#9 grouped by their enrichment in ASV_a45d (black) or ASV_65f4 (gray) is shown. Box plots were calculated with Statistical Analysis of Taxonomic and Functional Profiles (STAMP).

### Flagellar, ethanolamine, and galactose utilization genes were differentially enriched in the genomes related to ASV_65F4 and ASV_a45d

As mentioned, variants ASV_65f4 and ASV_a45d belong to different species of the genus *Flavonifractor*. The closest relative for ASV_65f4 was *F. plautii* (100% identical), while ASV_a45d was more closely related to *Flavonifractor* sp. strains An4 and An82 (98,6% identical), isolated by Medvecky and collaborators from the chicken cecum (38). These strains all have whole-genome assembly projects, thus, we performed a comparative genomic analysis to predict the gene functional differences between the taxonomic groups related to each variant. Specifically, we analyzed orthologous genes enriched in the group more closely related to ASV_65f4 (Table 1 and S6 Table) and more closely related to ASV_a45d (Table 2 and S7 Table). The ASV_65f4-related group was enriched in several genes involved in ethanolamine utilization (Table 1 and S6 Table). A closer examination revealed that both groups had ethanolamine utilization genes, but ASV_65f4-related group had these genes enriched, thus to establish the relevance of this enrichment, we reconstructed the operons for ethanolamine and propanediol utilization for both groups of genomes (propanediol catabolic pathway has homolog proteins to ethanolamine catabolic pathway that can be misannotated by automatic servers). The reconstruction of these operons revealed that two ethanolamine operons (Eut operon 1 and 2) are presented in the ASV_65f4-related group, while the ASV_a45d-related group only had Eut operon 1 (Fig 7). Eut operons 1 and 2 located in different parts of the genome and their proteins were highly similar but not identical (data not shown). Predicted proteins involved in the formation of flagella, flagellar proteins that interact with chemotaxis proteins, components of the flagellar motor that determine the direction of flagellar rotation, and the secretion of flagellar proteins were only presented in ASV_65f4-related group and *Flavonifractor* sp. strain An306 (Table 1 and S6 Table). The core set of flagellar genes (26 genes) (39) was identified in the 4 genomes belonging to ASV_65f4-related group, except for one of the genes that encode for a rod protein, FlgB, which was not found in *F. plautii* strain 2789STDY5834932. Meanwhile, the group more closely related to ASV_a45d was enriched in genes for galactose metabolism (Table 2 and S7 Table) and all had the complete pathway.

**Table 1.** Gene clusters enriched in the genomes more closely related to ASV_65f4. ASV_65f4-related group includes *F. plautii* YL31, *F. plautii* 2789STDY5834932, *F. plautii* ATCC 29863, and *F. plautii* An248.

| Cluster | Count | Name | p-value |
| --- | --- | --- | --- |
| Cluster 1 | 11 | ethanolamine catabolic process | 6.00E-07 |
| Cluster 2 | 4 | ornithine metabolic process | 0.00028198 |
| Cluster 3 | 3 | polyhedral organelle | 0.00255864 |
| Cluster 4 | 5 | bacterial-type flagellum-dependent swarming motility | 2.33E-08 |
| Cluster 5 | 5 | chemotaxis | 4.62E-07 |
| Cluster 6 | 4 | protein secretion | 3.91E-06 |
| Cluster 7 | 4 | bacterial-type flagellum-dependent cell motility | 1.14E-05 |

**Table 2.** Gene clusters enriched in the genomes more closely related to ASV_a45d. ASV_a45d-related group includes *Flavonifractor* sp. An4, *Flavonifractor* sp. An10, *Flavonifractor* sp. An82, *Flavonifractor* sp. An306.

| Cluster | Count | Name | p-value |
| --- | --- | --- | --- |
| Cluster 1 | 4 | Mo-molybdopterin cofactor biosynthetic process* | 8.54E-07 |
| Cluster 2 | 3 | riboflavin biosynthetic process* | 0.00044437 |
| Cluster 3 | 2 | galactose metabolic process | 0.00249661 |

\* Complete pathways for Riboflavin and Mo-molybdopterin cofactor biosynthetic were not present.

**Fig 7.**
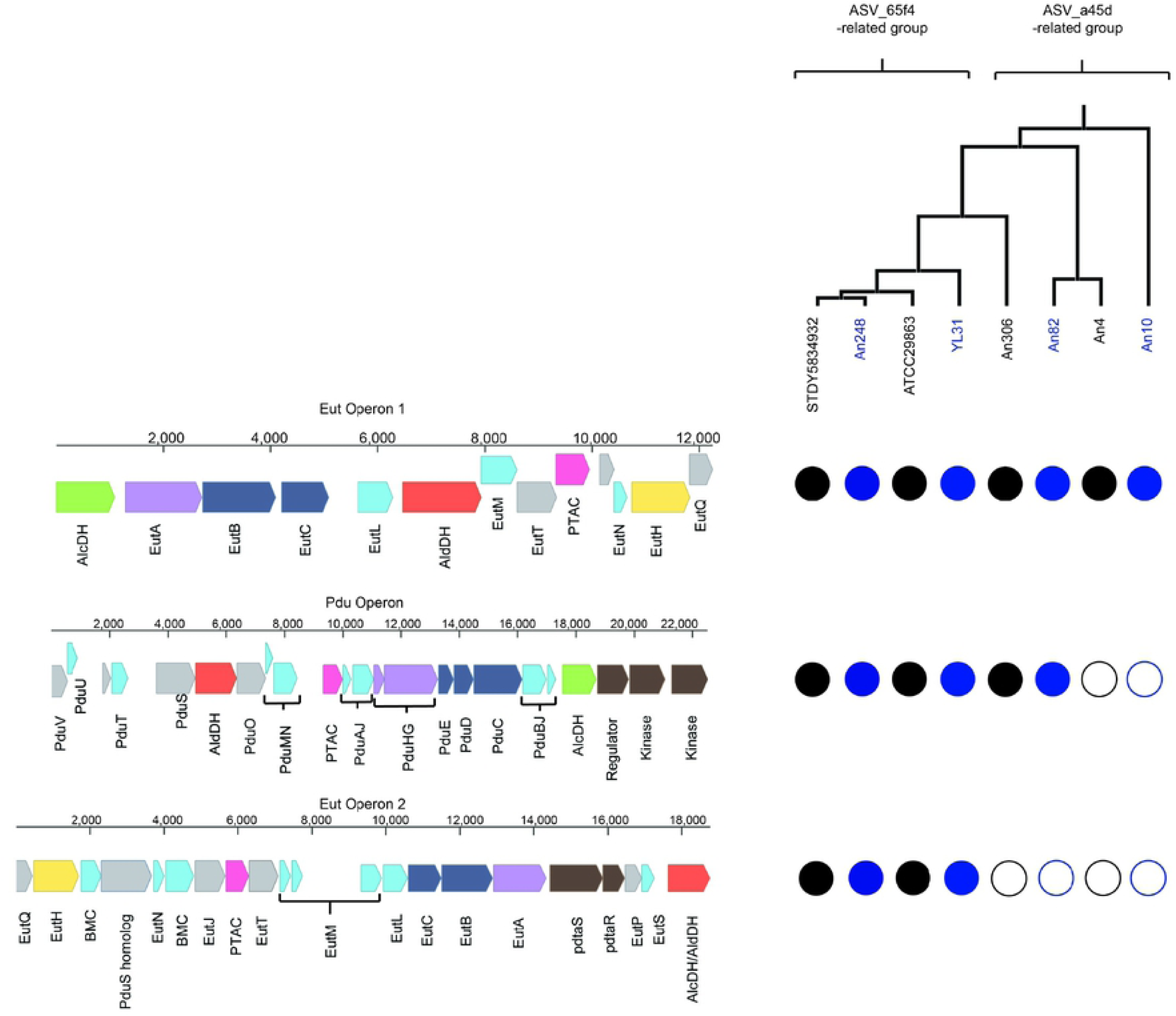
Representative BMC Loci for *Flavonifractor* spp. Cartoon representation of Ethanolamine utilization operon (Eut operon 1 and 2) and 1,2-propanediol utilization operon (Pdu operon). Genes are drawn on the *F. plautii* YL31 genome using Benchling [Biology Software] (2019). Eut operon 1 is 12,247 bp, Pdu operon is 22,527 bp, and Eut operon 2 is 18,769 bp. Abbreviations are as follows: AlcDH, Alcohol dehydrogenase; AldDH, Aldehyde dehydrogenase; PTAC, phosphotransacetylase; BMC, bacterial microcompartment; pdtaS, two-component system, sensor histidine kinase; pdtaR, two-component system, response regulator. Genes are color-coded according to their annotation: light blue, BMC-containing proteins; red, aldehyde dehydrogenase; green, alcohol dehydrogenase; solid pink, pduL-type phosphotransacylase; light purple, re-activating proteins; dark blue, signature enzymes (ethanolamine ammonia lyase subunits and propanediol dehydratase subunits); brown, regulatory element including two-component signaling elements; yellow, transporter; gray, other Eut or Pdu proteins. Circles show the presence (filled circle) or absence (white circle) of proteins in the strains depicted in the phylogenetic tree. The UPGMA phylogenetic tree involved 8 nucleotide sequences: *F. plautii* 2789STDY5834932 (STDY5834932), *F. plautii* An248 (An248), *F. plautii* ATCC 29863 (ATCC29863), *F. plautii* YL31 (YL31), *Flavonifractor* sp. An306 (An306), *Flavonifractor* sp. An82 (An82), *Flavonifractor* sp. An4 (An4), and *Flavonifractor* sp. An10 (An10). All positions containing gaps and missing data were eliminated. There were a total of 1008 positions in the final dataset. S8 Table contains all accession numbers for each protein in each strain. Pdu operon was reconstructed by its similarity with the one in *Salmonella enterica* subsp. *enterica* serovar Typhimurium str. LT2 (S9 Table).

### Predicted Phloretin Hydrolase is presented in both groups of genomes

Phloretin hydrolase gene (*phy*) is a well-characterized gene involved in the degradation of flavonoids. We found that both groups of genomes possess a protein similar to the reference Phloretin hydrolase (Table 3). Predicted Phy proteins of strains An4, An10, An82, and An306 were 90-95% identical to the reference Phy from *F. plautii* YL31.

**Table 3.**
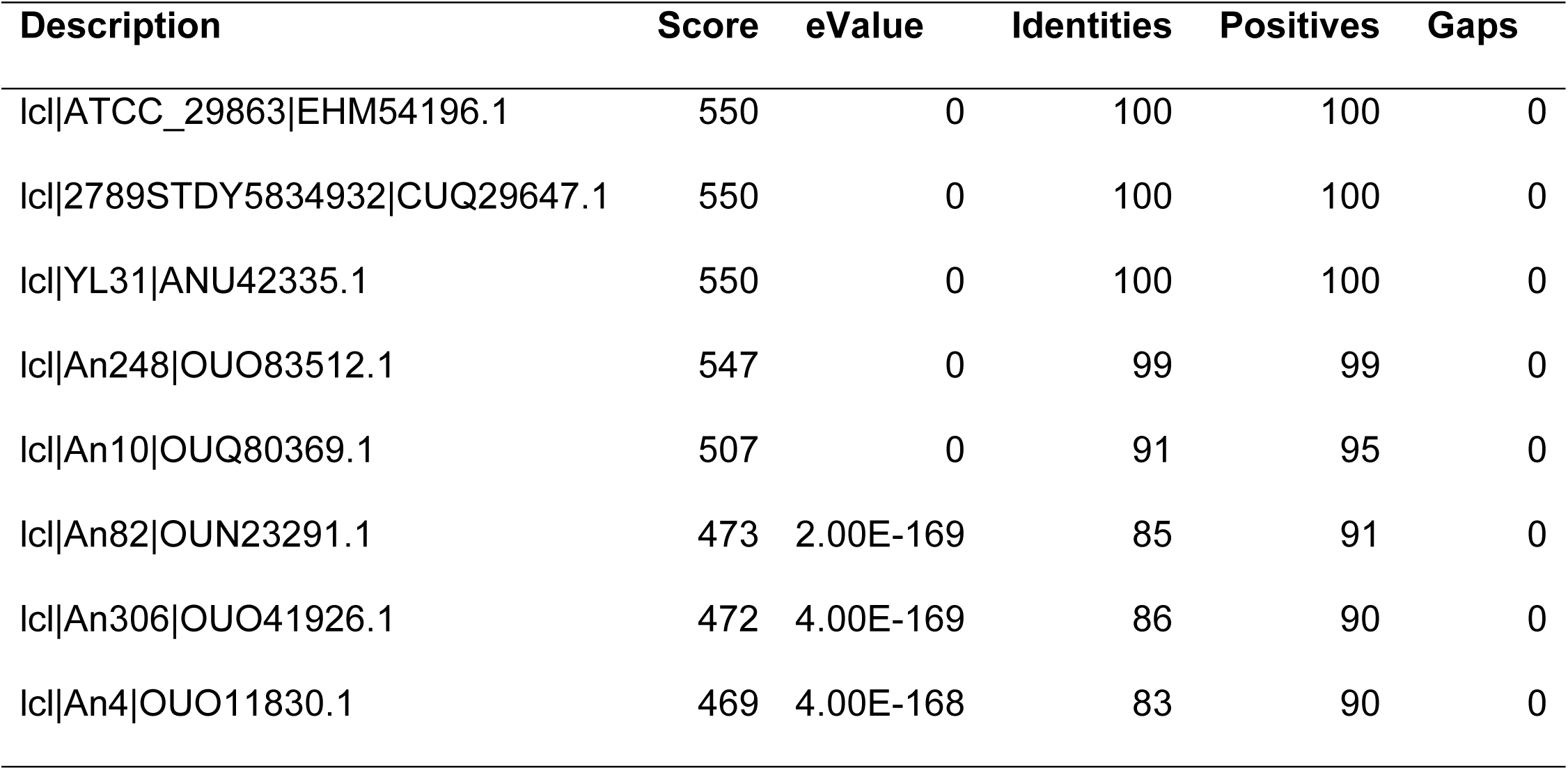
Blast results for Phloretin Hydrolase (Phy) for *Flavonifractor* spp.

## Discussion

Quercetin is presented in most fruits and its degradation produces biological active metabolites with effects on the host. Extending our knowledge of quercetin-degrading communities is important for predicting the health outcomes of flavonoid consumption. To study this matter, we used an *in vitro* fecal incubation system which can be used to directly evaluate the effect of the microbiota on a specific flavonoid. To limit the enrichment of non-quercetin degraders in these incubations, a medium low in carbon sources was used (20 mM of acetate). Under these conditions, mostly *Flavonifractor*-related sequences were enriched across libraries, specifically variants ASV_65f4 and ASV_a45d. These ASVs belong to different species, with 96.4 % of nucleotide identity in their 16S rRNA variable region V4 (40). The variant ASV_65f4 was 100 % identical to the known quercetin degrader *F. plautii* [42], which will explain its enrichment in the treatments. Meanwhile, the variant ASV_a45d was 98.6 % identical to *Flavonifractor* sp. An4 and An82 (38). The genomes of these species have a predicted phloretin hydrolase gene which catalyzes the hydrolytic C-C cleavage of phloretin, a flavonoid structurally similar to quercetin (41), generating phloroglucinol and 3-(4-hydroxyphenyl) propionic acid as biodegradation products. The enzyme is well characterized for another quercetin-degrader, *E. ramulus* (42). This indicates that strains An4 and An82 may harbor the enzymatic machinery necessary for the cleavage of the C-ring in quercetin as well, as *F. plautii* and *E. ramulus* do (43,44). In addition to the genome evidence of its close relatives, ASV_a45d was enriched in quercetin treatments with DOPAC production, even in the absence of other known quercetin degraders. Altogether this evidence suggests that ASV_a45d variant represents also a quercetin degrader.

ASVs related to *E. ramulus* were detected only in 4 out of 9 human fecal samples and none of the mice samples. However, *E. ramulus*-related ASVs were significantly increased by quercetin treatment only in one sample. *E. ramulus* was enriched in a previous study that supplemented with quercetin the diet of healthy volunteers under a flavonoid-free intervention (45). But under the culture conditions of this study, ASVs related to *Flavonifractor* were more prevalent. Another genus related to two of the ASVs enriched by quercetin treatments was *Intestinimonas*. Despite the relatedness of this bacterium with *Flavonifractor* sp., the ability to degrade quercetin was not detected in *Intestinimonas butyriciproducens* (46). However, it is not known whether this bacterium can use metabolites derived from quercetin degradation, like phloroglucinol. In some of our incubations, the genus *Coprococcus*, which is reported to use phloroglucinol (47), increased in relative abundance but did not reach significance.

Relative abundance correlations can be useful for identifying potential ecological interactions between microbial species (48). An interesting pattern of negative correlation was observed for *Flavonifractor*-related variants, ASV_65f4 and ASV_a45d. This may indicate a potential antagonistic interaction between these two species. Antagonism is more prevalent among phylogenetically and metabolically similar species. A study that screened 2,211 competing bacterial pairs from 8 different environments found that antagonism increased significantly between closely related strains and between strains that had a greater overlap in the capacity to grow on the 31 carbon sources screened through the Biolog assay (49). Since both ASVs from our study are phylogenetically related and both may have the capacity to degrade quercetin, this could explain the observed pattern of dominance of one or the other but not both. In competition assays, it is often the species that starts at high initial abundance the one that dominates (50). In our experiments, initial relative abundances did not predict which variant will dominate. Despite being less abundant at the initial point of incubation, ASV_a45d dominated over ASV_65f4 in half of the libraries after incubation with quercetin. However, when two fecal samples previously enriched in one or the other variant were combined, ASV_65f4 was always the strongest competitor and the dominant variant. Therefore, this evidence suggests that other processes besides initial relative abundances might be responsible for the dominance of the weaker competitor, ASV_a45d. Thus, through a comparative analysis of the genomes of close relatives, we sought functional capabilities that may allow one to thrive over the other.

*Flavonifractor* species clusters exhibited important differences in functional capacities. One clear difference found that might help explain why only ASV_65f4 increased in relative abundance in incubations supplemented with acetate but no quercetin was that ASV_65f4-related group had an enrichment of operons related to ethanolamine utilization. As far as we know, *Flavonifractor* sp. does not use acetate as a sole carbon source, thus, it might use ethanolamine under these culture conditions since this substrate can be generated from bacterial cell membranes. Ethanolamine can be used as carbon, nitrogen, and energy source by different bacteria (51). The reconstruction of the catabolic operon of this substrate in *Flavonifractor* spp. genomes revealed two ethanolamine operons (Eut operon 1 and 2), most similar to the reported EUT2 operon, which instead of the EutD phosphotransacetylase (PTAC) it encodes a homolog to the PduL PTAC, and in place of the EutR regulatory enzyme, it has a two-component regulatory system consisting of a signal transduction histidine kinase and a response regulator (52). Although both operons encode the essential proteins for ethanolamine utilization: EutBC protein, which is an ethanolamine ammonia lyase that converts ethanolamine into acetaldehyde and ammonia, the reactivating enzyme EutA that acts on EutBC, the aldehyde dehydrogenase (AldDH) EutE for converting acetaldehyde to acetyl-CoA that enters the tricarboxylic acid cycle and other processes, and the PduL PTAC homolog for the formation of acetate; Eut operon 2 had more *eut* genes that encode for structural proteins for the microcompartment (53). The catabolic process of ethanolamine occurs in microcompartments called metabolosomes, which are protein-based organelle-like structures that protect the cell from the potentially toxic effects of volatile intermediates; in the case of ethanolamine, acetaldehyde is this toxic intermediate. It is possible that harboring both operons (Eut operon 1 and 2) makes the group more closely related to ASV_65f4 more efficient for ethanolamine catabolism. The presence of ethanolamine catabolism in these strains might also give an advantage under nutrient scarcity in the gastrointestinal tract since this compound is abundant in this environment thanks to the action of phosphodiesterases on the phosphatidylethanolamine on bacterial and mammalian cell membranes which are constantly washed away in the mucus (54).

Another difference was observed in the carbohydrate metabolism of ASV_65f4- and ASV_a45d-related groups. The complete pathway for galactose utilization was only observed in the ASV_a45d-related group. Thus, these groups of strains might have an advantage under a diet based on dairy which is rich in lactose, a disaccharide formed from one molecule of glucose plus one of galactose. The two groups might harbor a structural difference too. The presence of a complete core set of flagellar genes was observed mostly in the ASV_65f4-related group. It has been reported that the *Flavonifractor* genus can be motile or non-motile (55). In the gastrointestinal tract, high levels of flagellin, a bacterial flagellar protein, have been associated with the breakdown of the intestinal barrier (56). We then suggest that the presence of flagellar genes and the utilization of ethanolamine make the ASV_65f4-related group more prone to opportunistic infections. Few infections by *F. plautii* have been reported (57,58).

The interactions of ASV_65f4 and ASV_a45d with other species in the microbial community might also explain the distinct enrichment of these bacteria if certain species favor one or the other variant. Identifying microbe-microbe co-occurrences is challenging because different samples might have a low number of shared species, obscuring the overall pattern of co-abundance that could identify interactions among groups. Despite this limitation, we were able to identify two genera that were associated with the dominance of ASV_65f4 or ASV_a45d. When ASV_65f4 dominated, the abundance of a succinate-utilizing bacterium, *Phascolarctobacterium*, was significantly higher. Two species of this genus are reported to only utilize succinate as a carbon source (59,60). Succinate is not a major fermentation product in human feces but saccharolytic bacteria that are abundant in the gastrointestinal tract can produce it and it is a substrate that bacteria can specialize in to coexist with bacteria that can readily utilize other more abundant carbon sources (59). Succinate is the product of fermentation of certain bacteria that use acetate as a carbon source and that under O_2_ depletion accumulate succinate, they slow down the tricarboxylic acid (TCA) cycle activity and shift to an alternative route called glyoxylate shunt-based TCA cycle that generates glyoxylate and succinate (61,62). The increase in abundance of *Phascolarctobacterium sp.* suggests that succinate became available, whether this is related to the presence of ASV_65f4 needs further experimental evidence.

Meanwhile, the relative abundance of *Desulfovibrio* sp. was significantly higher when ASV_a45d, the weakest competitor, became dominant. This abundance correlation could be explained by either a direct interaction between *Desulfovibrio* sp. and ASV_a45d or by an indirect effect, where *Desulfovibrio* sp. inhibits the stronger competitor, ASV_65f4. Evidence for the last argument could be related to the above-mentioned ethanolamine catabolism. *Desulfovibrio* sp. is an acetate utilizing bacterium that produces the toxic hydrogen sulfide (H_2_S) (63) (64). This metabolite has been observed to be involved in the ethanolamine metabolism of certain bacteria. During H_2_S detoxification, some bacteria convert it to thiosulfate, this has been observed in the gastrointestinal tract by the action of intestinal cells (65), and by bacteria in anoxic sediments (66,67). Bacteria can then convert thiosulfate to tetrathionate (68). When tetrathionate becomes available, it can serve as an electron donor for ethanolamine catabolism in some bacteria (69). Thus, in the presence of tetrathionate, more bacteria could consume ethanolamine increasing the competition for this substrate, which can influence ASV_65f4. Another mode of action of H_2_S on ethanolamine catabolism could be the inhibition of phosphodiesterases by H_2_S (70), which are the enzymes that convert phosphatidylethanolamine to ethanolamine. Other H_2_S-producing bacteria might affect ASV_65f4 as well, however, one of the reasons that *Desulfovibrio* sp. importance stood out in our experiments was that this genus was not affected by the diet fed to HMAM mice (data not shown). If the other variant, ASV_a45d, does not rely on ethanolamine for its carbon needs, this could explain why the presence of *Desulfovibrio* sp. only affects ASV_65f4.

The observations of this study show that *Flavonifractor*-related variants have the potential to utilize different carbon sources, interact with different species, and have different structural traits like motility. Thus, they might have a different impact on the host and the gut microbiome. Whether there is competition between *Flavonifractor*-related variants during flavonoid consumption warrants further investigation, as well as their metabolic capacity to degrade flavonoids, and the prevalence of these variants in other human populations.

## Acknowledgments

We would like to thank Dr. Robert Kerby (University of Wisconsin, Madison, USA) for his support and Dr. Bradley W. Bolling’s group (University of Wisconsin, Madison, USA) for their technical support processing the samples for HPLC analysis. We would like to thank the Doctorate program in Biosciences of La Sabana University and Ph.D. scholarship 647 (Colciencias, Colombia) for supporting Gina Paola Rodriguez.

## Author Contributions Statement

GR and FR conceived and planned the experiments. GR carried out the experiments. GR, AA, AC, and FR contributed to the interpretation of the results. AA, AC, and FR helped supervise the project and provided direction to the research. GR wrote the manuscript with input from all authors. All authors discussed the results and contributed to the final manuscript.

## Supporting information

**S1 Fig. Evolutionary relationships of *F. plautii* strains.**

**S2 Fig. Evolutionary relationships of *Flavonifractor* sp. strains.**

**S3 Fig. Genus abundance profiles from *in vitro* incubations with human fecal samples.**

**S4 Fig. Proportion of sequences (%) for ASV_65f4 in *in vitro* incubations with human fecal from subject #3.**

**S5 Fig. Univariate correlations between levels of six fecal taxa (ASVs) enriched in the presence of quercetin *in vitro*.**

**S6 Fig. Initial relative abundances for ASV_65f4 and ASV_a45d.**

**S1 Table. Genomes used in this study.**

**S2 Table. Average Nucleotide Identity (ANIm) and aligned percentage for 20 genomes belonging to *Flavonifractor* spp.**

**S3 Table. Concentration (mM) in culture of Quercetin and metabolites.**

**S4 Table. Statistical significance of Amplicon Sequence Variants (ASVs) enriched in quercetin treatments**.

**S5 Table. Estimates of Evolutionary Divergence between Amplicon Sequence Variants (ASVs) and Reference sequences.**

**S6 Table. Protein list for orthologous gene clusters enriched in ASV_65f4-related genomes**.

**S7 Table. Protein list for orthologous gene clusters enriched in ASV_a45d-related genomes**.

**S8 Table. Ids for Eut and Pdu proteins for *Flavonifractor* spp.**

**S9 Table. Distance Matrix for Pdu Operon in *F. plautii* YL31 with the one in *Salmonella enterica* subsp. *enterica* serovar Typhimurium str. LT2.**

